# *TREM2* is thyroid hormone regulated making the TREM2 pathway druggable with ligands for thyroid hormone receptor

**DOI:** 10.1101/2021.01.25.428149

**Authors:** Skylar J. Ferrara, Priya Chaudhary, Margaret J. DeBell, Gail Marracci, Hannah Miller, Evan Calkins, Edvinas Pocius, Brooke A. Napier, Ben Emery, Dennis Bourdette, Thomas S. Scanlan

**Affiliations:** Department of Chemical Physiology and Biochemistry and Program in Chemical Biology, Oregon Health & Science University, Portland, Oregon, USA; VA Portland Health Care System, Portland, Oregon, USA; Department of Biology, Portland State University, Oregon, USA; Department of Neurology, Oregon Health & Science University, Portland, Oregon, USA; Jungers Center for Neurosciences Research, Oregon Health & Science University, Portland, Oregon, USA

## Abstract

Triggering receptor expressed on myeloid cells-2 (TREM2) is a cell surface receptor on macrophages and microglia that senses and responds to disease associated signals to regulate the phenotype of these innate immune cells. The TREM2 signaling pathway has been implicated in a variety of diseases ranging from neurodegeneration in the central nervous system to metabolic disease in the periphery. We report here that *TREM2* is a thyroid hormone regulated gene and its expression in macrophages and microglia is stimulated by thyroid hormone. Both endogenous thyroid hormone and sobetirome, a synthetic thyroid hormone agonist drug, suppress pro-inflammatory cytokine production from myeloid cells including macrophages that have been treated with the SARS-CoV-2 spike protein which produces a strong, pro-inflammatory phenotype. Thyroid hormone agonism was also found to induce phagocytic behavior in microglia, a phenotype consistent with activation of the TREM2 pathway. The thyroid hormone antagonist NH-3 blocks the anti-inflammatory effects of thyroid hormone agonists and suppresses microglia phagocytosis. Finally, in a murine experimental autoimmune encephalomyelitis (EAE) multiple sclerosis model, treatment with Sob-AM2, a CNS-penetrating sobetirome prodrug, results in increased *Trem2* expression in disease lesion resident myeloid cells which correlates with therapeutic benefit in the EAE clinical score and reduced damage to myelin. Our findings represent the first report of endocrine regulation of *TREM2* and provide a unique opportunity to drug the TREM2 signaling pathway with orally active small molecule therapeutic agents.

## Introduction

Thyroid hormone (TH) provides essential regulation of many critical processes in vertebrate biology.^1^ Thyroxine (T4) is the predominant form of TH produced in and secreted from the thyroid gland, but its deiodinated metabolite 3,5,3’-triiodothyronine (T3) is the active form of TH that binds thyroid hormone receptors (TR) with high affinity. TRs bind to regulatory DNA sequences called thyroid hormone response elements (TRE) in the promoter regions of TH regulated genes, and T3 binding to TR in the cell nucleus activates TR to either stimulate or suppress transcription of these genes. Through such regulation of target genes TH action plays an important developmental role in the central nervous system (CNS) and periphery, as well as regulation of metabolism and homeostasis in many organs and cell types in the periphery. It is known that TH exerts effects on the immune system, in particular the innate immune cells, such as TR-expressing tissue-resident macrophages and microglia in the CNS; however, the mechanistic basis of TH-dependent effects on innate immunity is not well understood.^2^

Triggering receptor expressed on myeloid cells (TREM2) has emerged recently as a major regulator of the innate immune response and an important new therapeutic target connected to a number of diseases in the CNS and periphery.^3^ Expressed as a cell surface protein on macrophages and microglia, activation of TREM2 initiates a signal transduction cascade that triggers a switch in these cells away from a pro-inflammatory phenotype to an anti-inflammatory, phagocytic, restorative phenotype. Homozygous loss of function mutations in TREM2, or DAP12, a molecule that interacts with TREM2 to facilitate TREM2 signaling, causes Nasu-Hakola disease, a rare inborn error resulting in premature dementia, loss of myelin, and bone abnormalities.^4^ In addition, heterozygous TREM2 missense variants have been shown to be risk factors for common neurodegenerative diseases such as Alzheimer’s disease (AD), frontotemporal dementia, Parkinson’s disease (PD), and amyotrophic lateral sclerosis (ALS).^3, 5–6^ In the periphery, TREM2 expressed on circulating monocytes and tissue-resident macrophages has been implicated to play a beneficial role in controlling/resolving obesity/excess adiposity,^7^ non-alcoholic steatohepatitis (NASH),^8^ and hepatocellular carcinoma.^9^ In most of these diseases it appears that activation of the TREM2 pathway by increasing surface expression of TREM2 on macrophages and/or microglia and/or activating TREM2 with an agonist ligand is likely to be therapeutically beneficial. One of the main obstacles to targeting TREM2 for therapeutic benefit is that it falls into the category of “undruggable” or at least difficult to drug targets. This results from the nature of the ligands that bind to and activate TREM2. The general consensus is that TREM2 does not bind to a single, discrete ligand, but instead binds and is activated by various proteins, lipids, and other debris arising from damaged cells.^10^

Here we report the discovery that *Trem2* is a positively regulated thyroid hormone target gene. TREM2 expression and signaling through the TREM2 pathway is increased both *in vitro* and *in vivo* by treatment with T3, or the peripheral and CNS-penetrating thyromimetic agents sobetirome and Sob-AM2, respectively. This discovery effectively renders TREM2 “druggable” by systemically dosed small molecules with sufficient drug-like properties for targeting the TREM2 pathway in either the CNS or periphery.

## Results

Our interest in TREM2 was initiated from reports that microglia and macrophages increased phagocytic capacity upon treatment with the retinoid X receptor (RXR) agonist bexarotene,^11^ and treatment of TREM2-expressing myeloid lineage cells with other nuclear receptor agonists including PPAR and LXR also stimulated phagocytosis.^12^ Based on comprehensive genome-wide studies in mice, it has been reported that the *Trem2* gene is regulated by an RXR-dependent enhancer.^13^ Because a major biological function of RXR is to form heterodimers at positively-regulated response elements with thyroid hormone receptor (TR) and other members of nuclear receptor family including PPAR and LXR, we examined this enhancer from murine macrophages. We found the following sequence embedded in the putative RXR enhancer of *Trem2*—AGGGAG-GTTA-AGGTCA—which is a canonical DR4 thyroid hormone response element (TRE) (Fig.1). The AGGGAG is a consensus half-site for binding RXR, and AGGTCA is a consensus half-site for TR; the four intervening nucleotides (GTTA) make this a positively regulated DR4 TRE which should increase *TREM2* expression upon binding thyroid hormone agonists to TR (Fig. 1a).

**Figure 1:**
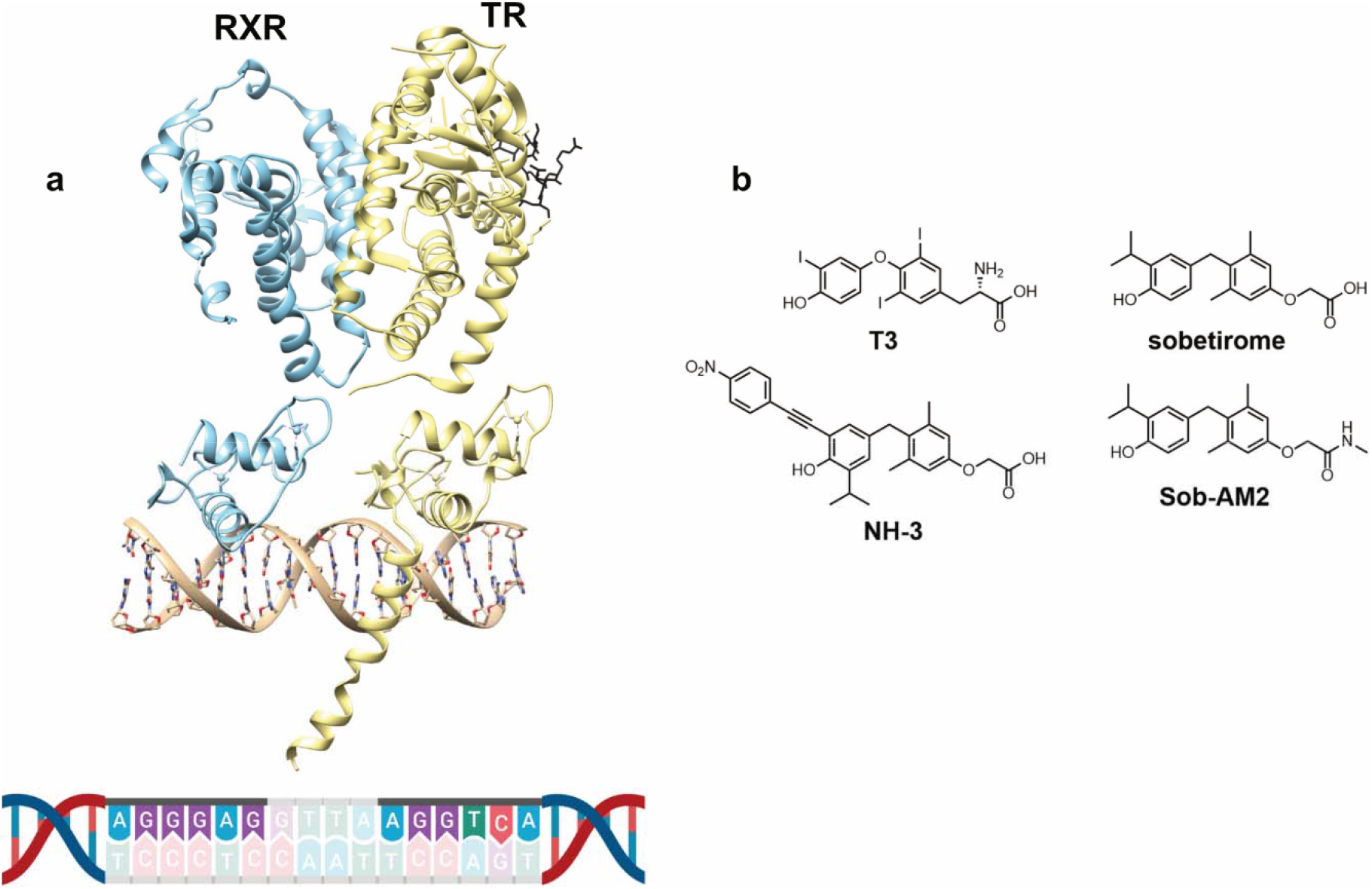
RXR-TR heterodimer associated with a DR4-TRE located in the promoter region of *TREM2* (**a**, PDB ID: 2NLL and 4ZO1) and endogenous and synthetic TR ligands used in this study (**b**).

We verified that *TREM2* expression is indeed positively regulated by thyroid hormone in both murine (Fig. 2a) and human (Fig. 2b) microglia, murine macrophages (Fig. 2c), and in mouse brain homogenate (Fig. 2d). Sobetirome is a synthetic T3 agonist drug that has a larger therapeutic index (TI) than T3, and Sob-AM2 is a prodrug of sobetirome that greatly facilitates delivery of sobetirome to the CNS from a systemic dose (Fig. 1b).^14–16^ We examined the ability of these agents to drive *Trem2* expression and found that sobetirome stimulates *Trem2* expression in mouse and human microglia (Fig. 2a,b), and hypothyroid mice treated with Sob-AM2 (i.p.) had increased *Trem2* expressed in brain compared to hypothyroid control (Fig. 2d). To verify the involvement of TR in the observed *TREM2* regulation by TH, the TR antagonist NH-3 (Fig. 1b) was employed in combination with T3.^17^ NH-3 was not only found to block T3 induction of *Trem2* expression in murine microglia, but to suppress *Trem2* expression significantly below the basal level observed in vehicle treated cells (Fig. 2a). As mentioned above, activation of the TREM2 signaling pathway induces both a pro-phagocytic and an anti-inflammatory response in monocytes and macrophages. We found that expression of the phagocytic marker *Cd68* was increased in microglia by treatment with either T3 or sobetirome, and the T3 effect was blocked by the TR antagonist NH-3 (Fig. 2e). In addition, expression of the pro-inflammatory cytokine interleukin-1β (*IL-1β*) was significantly decreased by T3 and sobetirome in mouse primary microglia (Fig. 2f).

**Figure 2:**
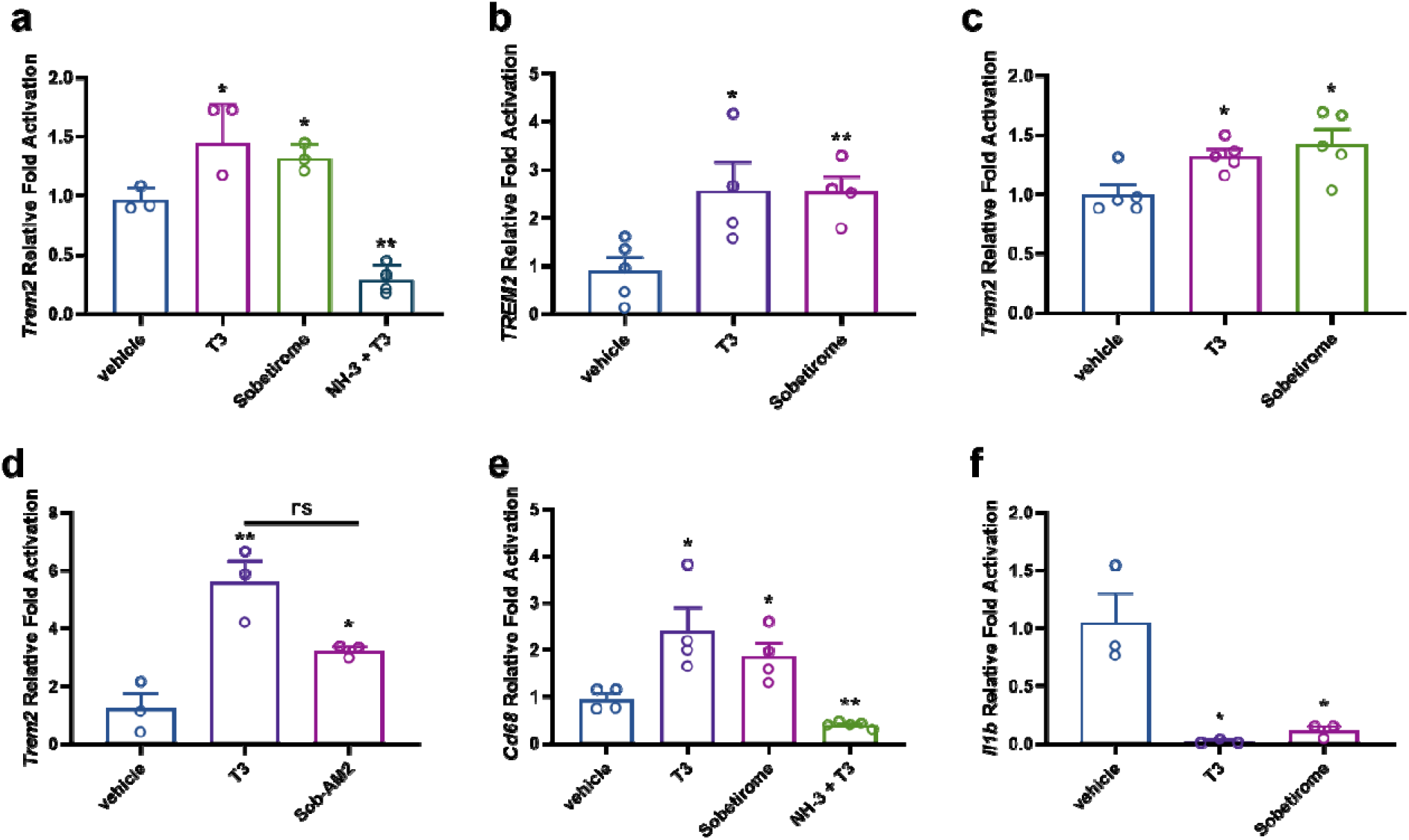
RT-qPCR demonstrating that T3 and thyromimetics regulate *TREM2* expression *in vitro* and *in vivo.* TR agonists T3 (10 nM) and sobetirome (1 μM) upregulate *TREM2* in mouse (**a**) and human (**b**) primary microglia, (**c**) mouse macrophage cell line RAW 264.7, **(d)** hypothyroid mouse whole brain extract (3.05 μmol/kg T3 and 30.5 μmol/kg Sob-AM2), and alter the transcript levels of TREM2 pathway-connected genes by upregulating (**e**) *Cd68* and downregulating (**f**) *Il1b* in mouse primary microglia (n = 3-5 as denoted). In contrast, TR antagonist NH-3 (2 μM NH-3 with 10 nM T3) downregulated *Trem2* (**a**) and *Cd68* (**e**) in mouse primary microglia. Statistical significance was determined by a 2-tailed, unpaired *t* test for comparisons between vehicle and group then were plotted together. Asterisks represent significant difference from vehicle unless otherwise noted (*P ≤ 0.05, **P ≤ 0.01, ***P ≤ 0.001). All graphs show mean ± SEM.

We next examined thyroid hormone regulated gene expression of the pro-inflammatory cytokine interleukin-6 (*IL-6*) to test the breadth of anti-inflammatory effects that would expected by thyromimetic stimulation of the TREM2 pathway. We tested whether *IL-6* expression stimulated from an inflammatory challenge in macrophages could be suppressed by thyromimetic stimulation of *TREM2* expression. (Fig. 3). It has been shown that macrophages incubated with coronavirus (SARS-CoV and SARS-CoV-2) spike S1 protein induce the production of pro-inflammatory cytokines including IL-6.^18–21^ RAW 264.7 cells stimulated with SARS-CoV-2 spike S1 protein induced a surge in *IL-6* expression compared to unstimulated cells, and this response was significantly attenuated upon treatment with T3 (Fig. 3a). T3 treatment also significantly suppressed *Il6* (Fig. 3b) and *Il1b* expression (Fig. 3c) in mouse primary lung macrophages stimulated with S1 protein. Concomitant with suppression of these pro-inflammatory cytokines, we observed stimulation of *Trem2* expression by T3 in primary lung macrophages (Fig. 3d).

**Figure 3:**
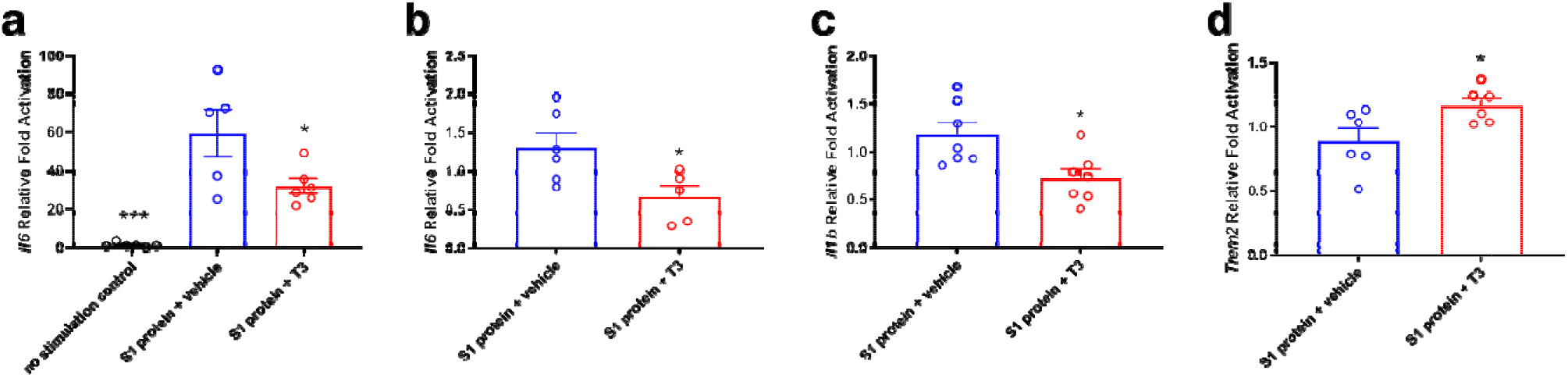
RT-qPCR of mouse RAW264.7 macrophage cells (**a**) and mouse primary lung macrophages (**b-d**) stimulated with 10 μg/mL SARS-CoV-2 S1 protein with and without treatment with 10 nM T3 (n = 5-7 as denoted). T3 treatment suppresses pro-inflammatory cytokine expression (*Il6* and *Il1b*, **a**-**c**) and upregulates *Trem2* expression following inflammatory stimulation with the S1 protein(**d**). Statistical significance was determined by a 2-tailed, unpaired *t* test for comparisons between vehicle and group then were plotted together. Asterisks represent significant difference from vehicle unless otherwise noted (*P ≤ 0.05, **P ≤ 0.01, ***P ≤ 0.001). All graphs show mean ± SEM.

The finding that TH agonists upregulate both *TREM2* and *CD68* expression predicts that phagocytosis will be stimulated by these drugs. This prompted us to directly examine phagocytosis, which is an established consequence of TREM2 signaling pathway activation in myeloid lineage cells. In order to ascertain the phagocytic response upon drug treatment, mouse primary microglia were cultured on glass coverslips and treated with drugs in the presence of fluorescent beads used as a phagocytosis substrate that can be monitored and quantified by fluorescence microscopy.^22^ Cellular bead uptake was evaluated via 3D volume visualization down orthogonal axes to confirm complete entrapment as shown in Fig. 4g. Microscopy of cells stained for CD11b microglial expression showed that T3 and sobetirome treatment significantly increased phagocytosis compared to vehicle control as judged by the number of beads engulfed by the microglia (Fig. 4a-c’). Conversely, the T3 antagonist NH-3 treatment was found to suppress phagocytic uptake of beads compared to vehicle-treated control (Fig. 4d,d’). Quantification of these microscopy data demonstrate that TH agonists stimulate, while TH antagonists block, phagocytosis in microglia (Fig. 4h). These results corroborate existing literature reporting augmented phagocytosis upon T3 treatment in macrophages.^23^

**Figure 4:**
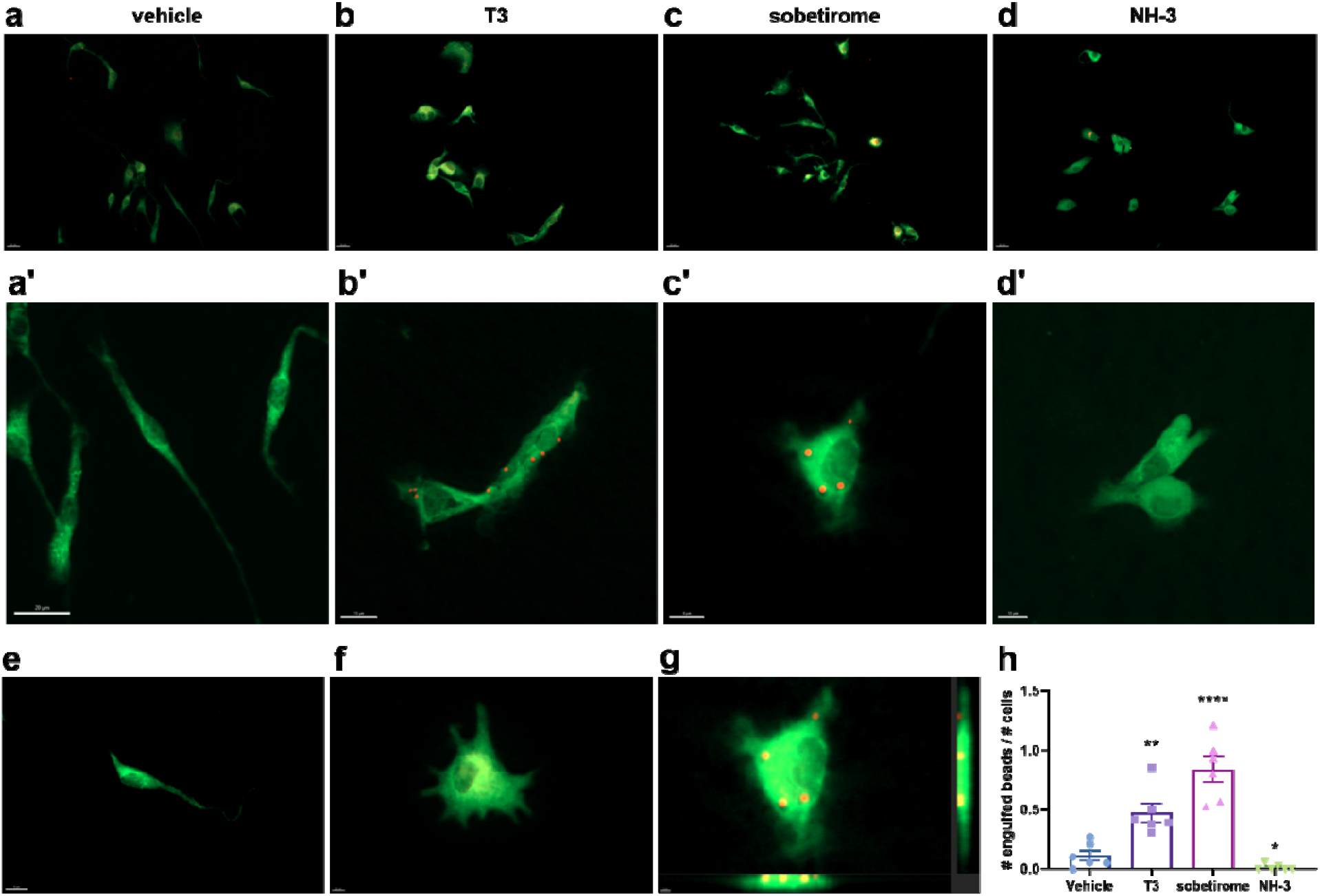
T3 stimulates and NH-3 blocks phagocytosis by microglia. C57BL/6 mouse primary microglia cells in culture were treated with DMSO vehicle (**a**+ **a’**), 10 nM T3 (**b**+ **b’**), 1 μM sobetirome (**c**+ **c’**), or 2 μM NH-3 (**d**+ **d’**) for 24 h before 2 μm diameter fluorescent beads (~100/cell) were introduced 2 h before cells were fixed and stained for Cd11b (green). (**e**) Elongated morphology of an unactivated/ramified microglia cell. (**f**) Morphology of an activated microglia cell upon treatment with sobetirome (see supporting information). (**g**) Three-dimensional view of beads inside the uppermost cell in the sobetirome treatment group (image generated in Imaris). (**h**) Quantification of phagocytosed beads per treatment group. Scale bars are 20 μm (**a-d**), 10-20 μm (**a’-d’**), and 8-10 μm (**e**-**g**) as noted. Statistical significance was determined by a 2-tailed, unpaired *t* test for comparisons between vehicle and group then were plotted together. Asterisks represent significant difference from vehicle unless otherwise noted (*P ≤ 0.05, **P ≤ 0.01, ***P ≤ 0.001). All graphs show mean ± SEM.

We have shown previously that CNS-penetrating thyromimetics stimulate myelin repair in different murine models of demyelination.^24^ One such model is experimental autoimmune encephalomyelitis (EAE), which is a demyelination model stimulated by an autoimmune attack that has parallels to the human disease multiple sclerosis. The autoimmune insult in EAE produces demyelination and axonal degeneration, particularly in the lumbar spinal cord region in mice. We have shown that in mice with EAE, T3, sobetirome, and Sob-AM2 treatment reduces clinical disease scores and demyelination and axonal degeneration within the spinal cord.^25^ Thus, the question arises as to whether this results in part from thyromimetic stimulation of the TREM2 signaling pathway in microglia and macrophages to induce phagocytosis of myelin debris, which is known to be a prerequisite to myelin repair.^26^ EAE was induced in C57Bl/6 mice within a 21-day treatment regimen, where mice were administered once daily injections of vehicle, T3, or Sob-AM2 starting 7 days after immunization before disease onset and continuing until day 21 when euthanasia and tissue collection occurred. Spinal cord sections co-stained with DAPI and TREM2 antibodies in the dorsal white matter of the lumbar section of spinal cords were analyzed for TREM2 content (Fig. 5a-c). Compared to vehicle, TREM2 protein expression was increased approximately four-fold and three-fold by T3 and Sob-AM2 treatment, respectively (Fig. 5d). Immunohistochemical analysis of this same spinal cord region showed that the population of CD11b positive cells, which correspond to microglia and/or macrophages, were not statistically different between Sob-AM2 treated mice and vehicle (see Supplementary Fig. S6).^25^ This indicates that the increase in TREM2 staining was not a result of increased myeloid cells in the spinal cord lesions. This increase in TREM2 protein expression observed by immunohistochemical analysis of spinal cord sections was confirmed by ELISA of spinal cord homogenate from unprocessed/unstained samples of the same groups (Fig. 5e). Improvement in EAE clinical scores (Fig. 5f) was observed in the mouse cohorts which were stained and correlates with augmented TREM2 expression.

**Figure 5:**
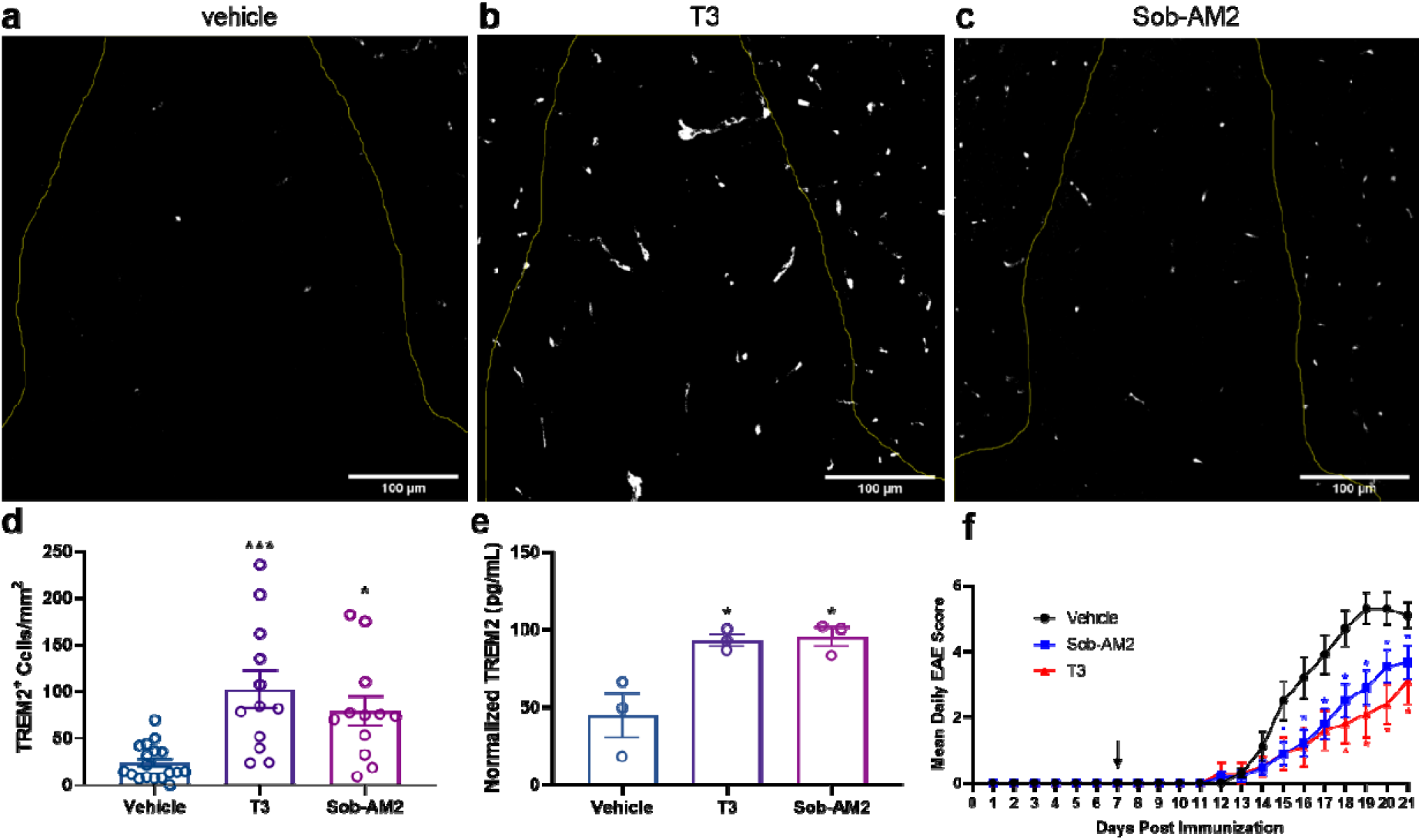
Treatment of EAE mice with T3 and Sob-AM2 upregulates *TREM2* expression in diseased spinal cord regions. **(a-c)** EAE mice were treated with vehicle, T3 (0.4 mg/kg), or Sob-AM2 (1 mg/kg) i.p. for 15 days (day 7-21). **(a-c)** Representative immunofluorescence images showing TREM2 protein expression within the dorsal white matter in the lumbar section of spinal cord. Scale bars: 100 μm. **(d)** Cells expressing TREM2 (white) within the dorsal white matter were quantified. T3 and Sob-AM2 increased TREM2 expression four-fold and three-fold compared to vehicle, respectively. Data represents images from vehicle (n = 18), T3 (n = 12), and Sob-AM2 (n = 12). Two sections were used from each mouse. Significance was determined using one-way ANOVA (P < 0.05). **(e)** ELISA measurement of TREM2 in spinal cord extracts from the same cohorts of EAE mice (n = 3) used in **a-d**. **(f)** Daily EAE clinical disease scores from vehicle (n = 18), T3 (0.4 mg/kg, n = 15), and Sob-AM2 (1 mg/kg, n = 28) treated cohorts. Statistical significance was determined by a two-tailed, unpaired *t* test comparing vehicle and treatment group per day with each *t* test performed independently. Asterisks represent significant difference from vehicle unless otherwise noted. (*P ≤ 0.05, **P ≤ 0.01, ***P ≤ 0.001). All graphs show mean ± SEM.

## Discussion

TREM2 has emerged as a central node in the innate immune response governing whether myeloid lineage macrophages and microglia have a pro-inflammatory or anti-inflammatory, restorative, healing phenotype in response to pathological insults. Here we report the discovery that *TREM2* is a direct target for regulation by thyroid hormone. The promoter region of the *TREM2* gene contains a consensus DR4-TRE which presents binding sites for the RXR-TR heterodimer that stimulates *TREM2* expression in response to thyroid hormone agonists. We found that *TREM2* expression was increased in both microglia and macrophages by addition of either T3 or the synthetic T3 analog sobetirome, and this T3 agonist induction of *TREM2* was blocked by the T3 antagonist NH-3. We also found that the proinflammatory cytokines IL-1β and IL-6 that are downregulated by induction of the TREM2 pathway were downregulated by T3 and sobetirome in microglia and macrophages that have been stimulated with the pro-inflammatory SARS-CoV-2 spike protein. Activation of the TREM2 pathway induces phagocytosis in these myeloid cells and T3 and sobetirome were found to induce phagocytosis in microglia whereas the T3 antagonist NH-3 blocked phagocytosis. Finally using the murine EAE model we showed that demyelinated regions of the spinal cord from mice treated with Sob-AM2, a CNS-penetrating prodrug of sobetirome, contained more TREM2 positive cells than control mice. Importantly, mice treated with Sob-AM2 had reduced clinical impairment, demyelination and axonal degeneration, suggesting that thyromimetic stimulation of TREM2 could result in protection in the inflammatory disease EAE. To our knowledge, ours is the first report that *TREM2,* and by extension the TREM2 pathway, is subject to endocrine regulation.

It has become increasingly apparent that thyroid hormone action plays a role in modulating innate immunity.^27^ Control of intracellular T3 levels in macrophages has been shown to be critical for these cells to respond appropriately to inflammatory signals.^28–29^ In addition, thyroid status dictates the degree to which rats respond to an inflammatory insult in which hypothyroidism induces increased pro-inflammatory cytokine production while hyperthyroidism inhibits this response.^30^ Administration of T3 blocks NLRP3 inflammasome activation, which depends upon NF-kB activation and results in IL-1β production and release, in different models of liver injury.^31–32^ In a model of kidney injury, deletion of the TRα isoforms result in increased injury, and isolated macrophages lacking TRα produce excessive levels of IL-1β compared to WT control.^33^ In addition to these examples of driving the transition to an anti-inflammatory phenotype in macrophages, TH has also been shown promote phagocytosis in macrophages,^23^ a response that we recapitulated here in microglia treated with T3. Collectively, these myeloid cell observations from both *in vitro* and *in vivo* experiments can be explained by thyroid hormone activation of the TREM2 pathway.

The sequelae of Nasu-Hakola disease provides genetic evidence that functional TREM2 and its signaling pathway in myeloid cells is essential for a normal, disease-free human lifespan.^4^ Over the past few years it has become increasingly apparent that the biology of the TREM2 signaling pathway is more involved in responding to pathological conditions as opposed to homeostatic physiology.^3^ For example, TREM2 is known to play a major role in a disparate collection of diseases including neurodegenerative diseases,^5, 34^ metabolic diseases,^7–8^ infectious diseases,^35–36^ cancer,^9, 37–39^ and stroke.^40^ Heterozygous missense mutations in *TREM2* are associated with strong increased risk of developing late onset Alzheimer’s disease (AD),^41–42^ and increased cerebrospinal fluid (CSF) levels of the soluble extracellular domain of TREM2 (sTREM2) has been shown to correlate with less disease severity in AD.^43^ This suggests that increased expression of TREM2 and/or increased signaling through the TREM2 pathway in microglia may be therapeutically beneficial in AD. It has also been known for some time that decreased thyroid hormone action in the CNS is detrimental while increased thyroid hormone action in the CNS correlates with clinical benefit in AD and other diseases of cognitive impairment.^44–50^ Our finding that TREM2 is a thyroid hormone regulated gene whose expression is increased by T3 acting on TR in microglia suggests a mechanistic connection between elevated levels of T3 in the CNS and clinical benefit in AD and related disorders. The beneficial effect of increased T3 in the brain may in part relate to increased stimulation of TREM2 expression in microglia and increased activity through the TREM2 pathway.

Another neurodegenerative disease that TREM2 appears associated with is multiple sclerosis (MS), an autoimmune disease that causes inflammatory demyelination and degeneration of axons in the CNS. Similar to AD, sTREM2 levels in the CSF are elevated in both relapsing remitting and primary progressive MS patients compared to CSF from patients with non-inflammatory neurological disease.^51^ In addition, TREM2-expressing myeloid cells have been detected in CSF and in demyelinating lesions upon autopsy from MS subjects.^51^ Macrophages and microglia release proinflammatory cytokines and probably contribute to demyelination and axonal degeneration in MS. In addition, microglia and macrophages play an important role in clearing myelin debris in demyelinating lesions that occur in MS via phagocytosis, which is a prerequisite to forming new myelin from mature oligodendrocytes.^26^ We have shown previously that the thyroid hormone receptor agonists sobetirome and Sob-AM2 promote remyelination in murine gliatoxin and genetic models of demyelination, and have further shown that like thyroid hormone, these agents stimulate oligodendrogenesis *in vitro* and *in vivo*.^24^ Our findings presented here that sobetirome induces TREM2 expression through TR activation in myeloid cells which stimulates phagocytosis in microglia constitutes a second beneficial mechanism of action in addition to oligodendrogenesis for thyroid hormone agonists as myelin repair agents.

This leads to the question of how the TREM2 pathway can be best engaged by drugs for therapeutic benefit. One approach would be to develop TREM2 agonists or antagonists that engage TREM2 directly to activate or block downstream signaling. A problem with this approach is that endogenous ligands for TREM2 appear to be a heterogenous collection of molecules associated with cell damage as opposed to a discrete metabolite or protein which is more amenable to drug discovery approaches. Some success has been reported with anti-TREM2 antibodies that act as agonists upon TREM2 binding,^52–53^ but the challenge with biologic agents such as antibodies is their intrinsic limits on distribution to different tissues and compartments affected by disease. For example, distribution from blood to the CNS is difficult with antibodies, and many of the diseases that would benefit from TREM2 engagement are localized in the CNS. Our findings that *TREM2* is a thyroid hormone regulated gene and that TREM2 expression and the TREM2 pathway can be activated or blocked by T3 agonists or antagonists opens the door to a therapeutic approach based on orally active small molecules. A number of studies have demonstrated that increased expression of TREM2 in microglia and/or macrophages alone serves to activate the TREM2 signaling pathway,^43, 54–56^ and this is consistent with our observations that T3 agonists increase TREM2 expression and activate the TREM2 pathway, whereas the T3 antagonist NH-3 blocks TREM2 expression and downstream signaling such as that involved in phagocytosis. The T3 agonist sobetirome would be representative of a drug candidate designed to stimulate the TREM2 pathway in the periphery for therapeutic benefit. Sobetirome therapy would be indicated, for example, for diseases of the liver that may benefit from activation of TREM2 such as NASH,^8^ immune-mediated damage following liver injury,^57^ and hepatocellular carcinoma.^9^ Alternatively, for peripheral diseases that may benefit from temporary blockade of the TREM2 pathway such as early in respiratory viral infection^35^ and certain cancers in which a robust immune response would be beneficial,^38–39^ the T3 antagonist NH-3 could be employed. However, during later stages of viral infection when cytokine storm can present, as can occur in COVID-19 critical illness, activation of the TREM2 pathway by sobetirome to drive macrophages to transition to an anti-inflammatory, resolving phenotype would be more appropriate for therapeutic benefit.^58–59^ The CNS-penetrating sobetirome prodrug Sob-AM2 would be the appropriate agent for AD, MS, and other neurodegenerative diseases for which therapeutic benefit would potentially come from TREM2 activation in the CNS while minimizing TREM2 activation and other thyromimetic activity in the periphery. And finally, should there be an application for blocking TREM2 activation in the CNS, the N-methyl amide prodrug of antagonist NH-3 provides enhanced distribution of NH-3 to the CNS while minimizing T3 antagonism in the periphery.^60^

In conclusion, we have found that *TREM2* is a direct target gene of thyroid hormone, making TREM2 and its signaling pathway in macrophages and microglia subject to regulation by a major endocrine system. To our knowledge ours is the first report of endogenous regulation of TREM2 at the level of gene expression, and it is likely to have important physiological and pathophysiological ramifications for TREM2-mediated innate immunity. This finding also represents a path toward developing small molecule therapeutics that either activate or suppress the TREM2 signaling pathway selectively in the CNS or periphery. The ability to “drug” TREM2 and its pathway with small molecule agents possessing good, drug-like properties would be an important medical advance for the diverse collection of diseases that intersect with TREM2 biology.

## Methods

### Reagents

T3 was purchased and used as received from Sigma-Aldrich. Sobetirome, Sob-AM2, and NH-3 were prepared as described in the literature.^14, 61–62^ Vehicle and drug stocks for cell culture experiments were prepared in dimethyl sulfoxide (DMSO). For *in vivo* experiments, all drugs were prepared at concentrations suitable for an i.p. injection of 150 μL per 26-g mouse. T3 drug stocks were prepared in 8 mM NaOH in saline and sobetirome and Sob-AM2 drug stocks were prepared in 50% DMSO in saline solutions. Vehicle stock solutions of 50% DMSO in saline (vehicle for sobetirome and Sob-AM2) or 8 mM NaOH in saline (T3 vehicle) were prepared and administered within the appropriate experiments. Drug concentrations are described in the figure legends for individual experiments. SARS-CoV-2 S1 protein was purchased from ACROBiosystems (#S1N-C52H3) and used as received. Information on other specific reagents is listed below within the Methods description for that particular experiment.

### Study approval and animals

All experiments were approved by the IACUC committee at the VA Portland Health Care System (VAPORHCS) or OHSU. Wild-type C57BL/6 mice, aged 8 to 10 weeks, were purchased from Jackson Laboratory and housed in climate-controlled rooms with a 12-hour light/12-hour dark cycle with *ad libitum* access to food and water. Mice were made hypothyroid by receiving 0.1% (w/v) methimazole and 0.2% (w/v) potassium perchlorate (Sigma-Aldrich) in drinking water for 2 weeks.^63^

### Cell culture

Mouse (ScienCell, #M1900-57) and human primary (Celprogen, #37089-01) microglia cells were purchased from and cultured according to the manufacturer’s protocol. RAW 264.7 cells were purchased fromm ATCC (ATCC^®^ TIB-71^™^) and cultured according to the manufacturer’s protocol in DMEM containing 10% FBS. Mouse primary lung macrophages were purchased from CellBiologics (#C57-2313F) and cultured according to the manufacturer’s protocol in DMEM containing 10% FBS. All cells were incubated at 37°C in the presence of 5% CO_2_. For experiments, cells were plated in either 6, 12, or 24 well plates at ~2 × 10^5^ cells per well.

### RT-qPCR

All cell cultures were serum-starved for 24 h before drug treatment, then treated with either DMSO vehicle, 10 nM T3, 1 μM sobetirome, or 2 μM NH-3 in the presence of 10 nM T3 for 24 h before RNA extraction. For mouse whole brain data, male C57Bl/6 mice (8-10 weeks old) were made hypothyroid according to standard procedure.^63^ Mouse cohorts were administered DMSO vehicle, 3.05 μmol/kg T3, or 30.5 μmol/kg Sob-AM2 and euthanized 6 h post-injection. Brains were immediately stored in 10-fold excess RNALater (Thermo Fisher) for RNA preservation. RNA was purified using either the RNeasy Mini Kit (Qiagen) or the PureLink RNA Mini kit with TRIzol extractions (Life Technologies) and quantified using a NanoDrop. PCR reactions were run on 1 μg of RNA per sample to afford cDNA using the QuantiTect Reverse Transcription kit (Qiagen). RT-qPCR was performed on an Applied Biosciences 7500 Real-Time PCR system following the QuantiTect SYBR Green PCR kit protocols (Qiagen) using Cyclophilin A (*Ppia*) and Glyceraldehyde 3-phosphate dehydrogenase (*GAPDH*) as housekeeping genes for mouse and human samples, respectively. Samples were run with technical duplicates and results were analyzed using the ΔΔCt relative quantification method.^64^

#### Mouse primers

f-*Ppia* 5′-AGGGTGGTGACTTTACACGC-3′, r-*Ppia* 5′-CTTGCCATCCAGCCATTCAG-3′; f-*Trem2* 5’-GACCTCTCCACCAGTTTCTCC-3’, r-*Trem2* 5’-TACATGACACCCTCAAGGACTG-3’; f-*Cd68* 5’-TTCTGCTGTGGAAATGCAAG-3’, r-*Cd68* 5’-GAGAAACATGGCCCGAAGT-3’; f-*Il1b* 5′-TCCAGGATGAGGACATGAGCAC-3′, r-*Il1b* 5′-GAACGTCACACACCAGCAGGTTA-3′, f-*Il6* 5’-GTTGCCTTCTTGGGACTGATG-3’, r-*Il6* 5’-CATACAATCAGAATTGCCATTGC-3’.^54, 65–68^

#### Human primers

f-*GAPDH* 5’-CAGGAGGCATTGCTGATGAT-3’, r-*GAPDH* 5’-GAAGGCTGGGGCTCATTT-3’; f-*TREM2* 5’-ACAGAAGCCAGGGACACATC-3’, r-*TREM2* 5’-CCTCCCATCATCTTCCTTCA-3’.^69–70^

##### Phagocytosis assay and immunocytochemistry

Mouse primary microglia cells were plated onto poly-L-lysine coated glass coverslips at ~5 × 10^4^ cells per coverslip seated within a 12 well plate (3 coverslips per group, 4 groups total). Cells were treated with DMSO vehicle, 10 nM T3, 1 μM sobetirome, or 2 μM NH-3 in the presence of 10 nM T3 for 24 h before the addition of 3 μL of a fluorescent latex bead suspension (L0280, Sigma) in complete DMEM in a ~100:1 bead-to-cell ratio for 2 h before the end of the experiment. Each well was then stripped of media, washed with PBS, and cells were fixed with 4% paraformaldehyde in PBS for 10 min at room temperature. The cells were then permeabilized with PBS containing 0.05% saponin (saponin was included in all subsequent incubations and washes) and stained for microglial marker CD11b using monoclonal rat anti-mouse CD11b (1:200 dilution, AbD Serotec, #MCA711) in conjunction with Alexa Fluor 488-conjugated donkey anti-mouse secondary antibody (1:400 dilution, Thermo Fisher, A21202). Coverslips were then washed and mounted with ProLong^®^ Gold antifade reagent (Life Technologies). Two fields per coverslip were imaged for a total of six fields of microglial cells per group imaged with a Zeiss ApoTome.2 at 20x magnification collecting z-stack images to verify the entrapment of fluorescent beads within the CD11b-stained cell field. Images were acquired and processed with ZEN 2 (blue edition) version 3.1 software (Zeiss), ImageJ/FIJI, and IMARIS software (Bitplane). Blue fluorescent beads were colorized to red during image post-processing in IMARIS for ease of visualization.

### EAE experiment, histology, and immunofluorescence

Immunization of female C57BL/6 mice (The Jackson Laboratory, Bar Harbor, ME, ages 8-10 weeks) with 200 μg of myelin oligodendrocyte glycoprotein (MOG) 35-55 (PolyPeptide Laboratories, San Diego, CA) in complete Freund’s adjuvant containing 400 μg of *Mycobacterium tuberculosis* per mouse (subcutaneous injection of 0.2ml volume), followed by pertussis toxin (List Biological labs Inc), was administered via intraperitoneal (i.p.) injection at day 0 (75ng per mouse) and day 2 (200ng per mouse) after immunization. All mice were scored for clinical signs of EAE daily using a 9-point scale and received one-time daily i.p. injections of vehicle (50% DMSO or 8mM NaOH, both in saline) or drug (T3 0.4 mg/kg or Sob-AM2 1 mg/kg) starting at day 7 through euthanasia on day 21 post-immunization. Each group contained 6-8 mice and the experiment was repeated 3 times. At 21 days post-immunization, mice were euthanized with carbon dioxide and spinal columns were removed. Columns were immersed in 4% paraformaldehyde (PFA) for 24-48 hours, then spinal cords were extracted and fixed in a microwave for 1 hour. Free-floating 40 μm lateral sections were collected from the lumbar region using a vibratome, then stored in PBS at 4°C. Tissues were permeabilized with 0.05% Triton X-100 in PBS for 30 minutes, washed in PBS, then blocked with 5% donkey serum in PBS for 3 hours. Sections were incubated in primary monoclonal anti-TREM2 antibody (EMD Millore, MABN2320, 1:250) in blocking buffer overnight at 4°C. Tissues were washed in PBS, then incubated in Alexa Fluor 647-conjugated goat anti-rat secondary antibody (Thermo Fisher, A21247, 1:200) and DAPI (1:50000) in PBS overnight at 4°C. Sections were washed and mounted on slides using ProLong^®^ Gold antifade reagent (Life Technologies) and imaged using Zeiss 780 laser scanning confocal microscope at 20x. Cells expressing TREM2 were quantified within the region of dorsal white matter. Images were acquired and processed with ZEN 2 (blue edition) version 3.1 software (Zeiss), ImageJ/FIJI, and IMARIS software (Bitplane). TREM2 was colorized white in ImageJ/FIJI during post-processing for ease of visualization. Quantification of TREM2 concentration via ELISA was performed using a TREM2 ELISA kit (Reddot Biotech Inc). following the manufacturer’s instructions.

### Statistical analysis

Statistical significance was determined using 1-way ANOVA with Dunnett’s post-test or two-tailed, unpaired Student’s *t* tests between two groups and then plotted together graphically as denoted in each figure legend (P < 0.05). For *in vivo* experiments, there were no differences in effects between the different vehicles (8 mM NaOH in saline or 50% DMSO in saline), so data from the different vehicles were combined into a single vehicle control group. Replicates for each experiment were as stated in the specific figure legend and in the corresponding methods. Sample sizes for animal experiments were informed by previous literature accounts or from preliminary data to minimize total animal numbers as appropriate. Analysis was carried out in GraphPad Prism 8 without further modifications. The ROUT method was used to identify and eliminate outliers. Significance level was set to \**P* ≤ 0.05, \*\**P* ≤ 0.01, \*\*\**P* ≤ 0.001, and \*\*\*\**P* ≤ 0.0001. All graphs show mean ± SEM.

## Author contributions

S.J.F. and T.S.S. conceived of the project and experiments. S.J.F. and H.M. performed cell culture experiments and RT-qPCR. S.J.F. performed the phagocytosis assay and associated immunocytochemistry and microscopy. G.M., E.P., E.C., and P.C. conducted the EAE experiment and tissue procurement. S.J.F. and M.J.D. stained spinal cords and performed the associated microscopy and data analysis. S.J.F. synthesized the relevant compounds and performed overall data analysis. B.E., B.A.N., and D.B. provided advice on experimental design. S.J.F. and T.S.S. wrote the manuscript with input from P.C., B.E., B.A.N., and D.B.

## Acknowledgments

This research was supported by NIH grants DK52798 (T.S.S.) and GM133804 (B.A.N.), and the National Multiple Sclerosis Society grants RG 5199A4 and RG-1607-25053 to D.B., RG 5106A1/1 and RG-2001-35775 to B.E., the Race to Erase MS to D.B., the OHSU Laura Fund for Innovation in Multiple Sclerosis to D.B. and T.S.S. We would like to thank the Advanced Light Microscopy Core (supported by NIH P30 NS061800) at OHSU for technical assistance.

## References

1. Yen, P. M., Physiological and Molecular Basis of Thyroid Hormone Action. Physiological Reviews 2001, 81 (3), 1097–1142.

2. De Vito, P.; Incerpi, S.; Pedersen, J. Z.; Luly, P.; Davis, F. B.; Davis, P. J., Thyroid Hormones as Modulators of Immune Activities at the Cellular Level. Thyroid 2011, 21 (8), 879–890.

3. Deczkowska, A.; Weiner, A.; Amit, I., The Physiology, Pathology, and Potential Therapeutic Applications of the TREM2 Signaling Pathway. Cell 2020, 181 (6), 1207–1217.

4. Xing, J.; Titus, A. R.; Humphrey, M. B., The TREM2-DAP12 signaling pathway in Nasu-Hakola disease: a molecular genetics perspective. Res Rep Biochem 2015, 5, 89–100.

5. Krasemann, S.; Madore, C.; Cialic, R.; Baufeld, C.; Calcagno, N.; El Fatimy, R.; Beckers, L.; O’Loughlin, E.; Xu, Y.; Fanek, Z.; Greco, D. J.; Smith, S. T.; Tweet, G.; Humulock, Z.; Zrzavy, T.; Conde-Sanroman, P.; Gacias, M.; Weng, Z.; Chen, H.; Tjon, E.; Mazaheri, F.; Hartmann, K.; Madi, A.; Ulrich, J. D.; Glatzel, M.; Worthmann, A.; Heeren, J.; Budnik, B.; Lemere, C.; Ikezu, T.; Heppner, F. L.; Litvak, V.; Holtzman, D. M.; Lassmann, H.; Weiner, H. L.; Ochando, J.; Haass, C.; Butovsky, O., The TREM2-APOE Pathway Drives the Transcriptional Phenotype of Dysfunctional Microglia in Neurodegenerative Diseases. Immunity 2017, 47 (3), 566–581.e9.

6. Ulrich, J. D.; Holtzman, D. M., TREM2 Function in Alzheimer’s Disease and Neurodegeneration. ACS Chemical Neuroscience 2016, 7 (4), 420–427.

7. Jaitin, D. A.; Adlung, L.; Thaiss, C. A.; Weiner, A.; Li, B.; Descamps, H.; Lundgren, P.; Bleriot, C.; Liu, Z.; Deczkowska, A.; Keren-Shaul, H.; David, E.; Zmora, N.; Eldar, S. M.; Lubezky, N.; Shibolet, O.; Hill, D. A.; Lazar, M. A.; Colonna, M.; Ginhoux, F.; Shapiro, H.; Elinav, E.; Amit, I., Lipid-Associated Macrophages Control Metabolic Homeostasis in a Trem2-Dependent Manner. Cell 2019, 178 (3), 686–698.e14.

8. Xiong, X.; Kuang, H.; Ansari, S.; Liu, T.; Gong, J.; Wang, S.; Zhao, X.-Y.; Ji, Y.; Li, C.; Guo, L.; Zhou, L.; Chen, Z.; Leon-Mimila, P.; Chung, M. T.; Kurabayashi, K.; Opp, J.; Campos-Pérez, F.; Villamil-Ramírez, H.; Canizales-Quinteros, S.; Lyons, R.; Lumeng, C. N.; Zhou, B.; Qi, L.; Huertas-Vazquez, A.; Lusis, A. J.; Xu, X. Z. S.; Li, S.; Yu, Y.; Li, J. Z.; Lin, J. D., Landscape of Intercellular Crosstalk in Healthy and NASH Liver Revealed by Single-Cell Secretome Gene Analysis. Molecular Cell 2019, 75 (3), 644–660.e5.

9. Tang, W.; Lv, B.; Yang, B.; Chen, Y.; Yuan, F.; Ma, L.; Chen, S.; Zhang, S.; Xia, J., TREM2 acts as a tumor suppressor in hepatocellular carcinoma by targeting the PI3K/Akt/β-catenin pathway. Oncogenesis 2019, 8 (2), 9.

10. Kober, D. L.; Brett, T. J., TREM2-Ligand Interactions in Health and Disease. Journal of Molecular Biology 2017, 429 (11), 1607–1629.

11. Natrajan, M. S.; de la Fuente, A. G.; Crawford, A. H.; Linehan, E.; Nuñez, V.; Johnson, K. R.; Wu, T.; Fitzgerald, D. C.; Ricote, M.; Bielekova, B.; Franklin, R. J. M., Retinoid X receptor activation reverses age-related deficiencies in myelin debris phagocytosis and remyelination. Brain 2015, 138 (12), 3581–3597.

12. Savage, J. C.; Jay, T.; Goduni, E.; Quigley, C.; Mariani, M. M.; Malm, T.; Ransohoff, R. M.; Lamb, B. T.; Landreth, G. E., Nuclear Receptors License Phagocytosis by Trem2^+^ Myeloid Cells in Mouse Models of Alzheimer’s Disease. The Journal of Neuroscience 2015, 35 (16), 6532–6543.

13. Daniel, B.; Nagy, G.; Hah, N.; Horvath, A.; Czimmerer, Z.; Poliska, S.; Gyuris, T.; Keirsse, J.; Gysemans, C.; Van Ginderachter, J. A.; Balint, B. L.; Evans, R. M.; Barta, E.; Nagy, L., The active enhancer network operated by liganded RXR supports angiogenic activity in macrophages. Genes & Development 2014, 28 (14), 1562–1577.

14. Meinig, J. M.; Ferrara, S. J.; Banerji, T.; Banerji, T.; Sanford-Crane, H. S.; Bourdette, D.; Scanlan, T. S., Targeting Fatty-Acid Amide Hydrolase with Prodrugs for CNS-Selective Therapy. ACS Chemical Neuroscience 2017, 8 (11), 2468–2476.

15. Meinig, J. M.; Ferrara, S. J.; Banerji, T.; Banerji, T.; Sanford-Crane, H. S.; Bourdette, D.; Scanlan, T. S., Structure–Activity Relationships of Central Nervous System Penetration by Fatty Acid Amide Hydrolase (FAAH)-Targeted Thyromimetic Prodrugs. ACS Medicinal Chemistry Letters 2019, 10 (1), 111–116.

16. Scanlan, T. S., Sobetirome: a case history of bench-to-clinic drug discovery and development. Heart Failure Reviews 2010, 15 (2), 177–182.

17. Nguyen, N.-H.; Apriletti, J. W.; Cunha Lima, S. T.; Webb, P.; Baxter, J. D.; Scanlan, T. S., Rational Design and Synthesis of a Novel Thyroid Hormone Antagonist That Blocks Coactivator Recruitment. Journal of Medicinal Chemistry 2002, 45 (15), 3310–3320.

18. Wang, W.; Ye, L.; Ye, L.; Li, B.; Gao, B.; Zeng, Y.; Kong, L.; Fang, X.; Zheng, H.; Wu, Z.; She, Y., Up-regulation of IL-6 and TNF-α induced by SARS-coronavirus spike protein in murine macrophages via NF-κB pathway. Virus Research 2007, 128 (1), 1–8.

19. Dosch, S. F.; Mahajan, S. D.; Collins, A. R., SARS coronavirus spike protein-induced innate immune response occurs via activation of the NF-κB pathway in human monocyte macrophages in vitro. Virus Research 2009, 142 (1), 19–27.

20. chen, y.; Feng, Z.; Diao, B.; Wang, R.; Wang, G.; Wang, C.; Tan, Y.; Liu, L.; Wang, C.; Liu, Y.; Liu, Y.; Yuan, Z.; Ren, L.; Wu, Y., The Novel Severe Acute Respiratory Syndrome Coronavirus 2 (SARS-CoV-2) Directly Decimates Human Spleens and Lymph Nodes. medRxiv 2020, 2020.03.27.20045427.

21. Theobald, S.; Simonis, A.; Kreer, C.; Zehner, M.; Fischer, J.; Albert, M.-C.; Malin, J.; Gräb, J.; Winter, S.; Silva, U. S. d.; Böll, B.; Köhler, P.; Gruell, H.; Suàrez, I.; Hallek, M.; Fätkenheuer, G.; Jung, N.; Cornely, O.; Lehmann, C.; Kashkar, H.; Klein, F.; Rybniker, J., The SARS-CoV-2 spike protein primes inflammasome-mediated interleukin-1-beta secretion in COVID-19 patient-derived macrophages. Research Square: 2020.

22. Lian, H.; Roy, E.; Zheng, H., Microglial Phagocytosis Assay. Bio-protocol 2016, 6 (21), e1988.

23. Perrotta, C.; Buldorini, M.; Assi, E.; Cazzato, D.; De Palma, C.; Clementi, E.; Cervia, D., The Thyroid Hormone Triiodothyronine Controls Macrophage Maturation and Functions: Protective Role during Inflammation. The American Journal of Pathology 2014, 184 (1), 230–247.

24. Hartley, M. D.; Banerji, T.; Tagge, I. J.; Kirkemo, L. L.; Chaudhary, P.; Calkins, E.; Galipeau, D.; Shokat, M. D.; DeBell, M. J.; Van Leuven, S.; Miller, H.; Marracci, G.; Pocius, E.; Banerji, T.; Ferrara, S. J.; Meinig, J. M.; Emery, B.; Bourdette, D.; Scanlan, T. S., Myelin repair stimulated by CNS-selective thyroid hormone action. JCI Insight 2019, 4 (8).

25. Chaudhary, P.; Marracci, G. H.; Calkins, E.; Pocius, E.; Bensen, A. L.; Scanlan, T. S.; Emery, B.; Bourdette, D. N., Thyroid hormone and thyromimetics inhibit myelin and axonal degeneration and oligodendrocyte loss in EAE. bioRxiv 2020, 2020.12.20.423638.

26. Lloyd, A. F.; Miron, V. E., The pro-remyelination properties of microglia in the central nervous system. Nature Reviews Neurology 2019, 15 (8), 447–458.

27. Montesinos, M. d. M.; Pellizas, C. G., Thyroid Hormone Action on Innate Immunity. Frontiers in Endocrinology 2019, 10 (350).

28. van der Spek, A. H.; Surovtseva, O. V.; Jim, K. K.; van Oudenaren, A.; Brouwer, M. C.; Vandenbroucke-Grauls, C. M. J. E.; Leenen, P. J. M.; van de Beek, D.; Hernandez, A.; Fliers, E.; Boelen, A., Regulation of Intracellular Triiodothyronine Is Essential for Optimal Macrophage Function. Endocrinology 2018, 159 (5), 2241–2252.

29. Anne, H. v. d. S.; Eric, F.; Anita, B., Thyroid hormone metabolism in innate immune cells. Journal of Endocrinology 2017, 232 (2), R67–R81.

30. Rittenhouse, P. A.; Redei, E., Thyroxine Administration Prevents Streptococcal Cell Wall-Induced Inflammatory Responses*. Endocrinology 1997, 138 (4), 1434–1439.

31. Vargas, R.; Videla, L. A., Thyroid hormone suppresses ischemia-reperfusion-induced liver NLRP3 inflammasome activation: Role of AMP-activated protein kinase. Immunology Letters 2017, 184, 92–97.

32. Dong, X.; Yang, H.; Li, C.; Liu, Q.; Bai, Q.; Zhang, Z., Triiodothyronine alleviates alcoholic liver disease injury through the negative regulation of the NLRP3 signaling pathway. Exp Ther Med 2018, 16 (3), 1866–1872.

33. Furuya, F.; Ishii, T.; Tamura, S.; Takahashi, K.; Kobayashi, H.; Ichijo, M.; Takizawa, S.; Kaneshige, M.; Suzuki-Inoue, K.; Kitamura, K., The ligand-bound thyroid hormone receptor in macrophages ameliorates kidney injury via inhibition of nuclear factor-κB activities. Scientific Reports 2017, 7 (1), 43960.

34. Konishi, H.; Kiyama, H., Microglial TREM2/DAP12 Signaling: A Double-Edged Sword in Neural Diseases. Frontiers in Cellular Neuroscience 2018, 12 (206).

35. Wu, K.; Byers, D. E.; Jin, X.; Agapov, E.; Alexander-Brett, J.; Patel, A. C.; Cella, M.; Gilfilan, S.; Colonna, M.; Kober, D. L.; Brett, T. J.; Holtzman, M. J., TREM-2 promotes macrophage survival and lung disease after respiratory viral infection. Journal of Experimental Medicine 2015, 212 (5), 681–697.

36. Chen, Q.; Zhang, K.; Jin, Y.; Zhu, T.; Cheng, B.; Shu, Q.; Fang, X., Triggering receptor expressed on myeloid cells-2 protects against polymicrobial sepsis by enhancing bacterial clearance. American journal of respiratory and critical care medicine 2013, 188 (2), 201–12.

37. Donatelli, S. S.; Zhou, J. M.; Gilvary, D. L.; Eksioglu, E. A.; Chen, X.; Cress, W. D.; Haura, E. B.; Schabath, M. B.; Coppola, D.; Wei, S.; Djeu, J. Y., TGF-β-inducible microRNA-183 silences tumor-associated natural killer cells. Proceedings of the National Academy of Sciences of the United States of America 2014, 111 (11), 4203–8.

38. Yao, Y.; Li, H.; Chen, J.; Xu, W.; Yang, G.; Bao, Z.; Xia, D.; Lu, G.; Hu, S.; Zhou, J., TREM-2 serves as a negative immune regulator through Syk pathway in an IL-10 dependent manner in lung cancer. Oncotarget 2016, 7 (20), 29620–29634.

39. Zhang, X.; Wang, W.; Li, P.; Wang, X.; Ni, K., High TREM2 expression correlates with poor prognosis in gastric cancer. Human pathology 2018, 72, 91–99.

40. Gervois, P.; Lambrichts, I., The Emerging Role of Triggering Receptor Expressed on Myeloid Cells 2 as a Target for Immunomodulation in Ischemic Stroke. Frontiers in immunology 2019, 10, 1668.

41. Guerreiro, R.; Wojtas, A.; Bras, J.; Carrasquillo, M.; Rogaeva, E.; Majounie, E.; Cruchaga, C.; Sassi, C.; Kauwe, J. S. K.; Younkin, S.; Hazrati, L.; Collinge, J.; Pocock, J.; Lashley, T.; Williams, J.; Lambert, J.-C.; Amouyel, P.; Goate, A.; Rademakers, R.; Morgan, K.; Powell, J.; St. George-Hyslop, P.; Singleton, A.; Hardy, J., TREM2 Variants in Alzheimer’s Disease. New England Journal of Medicine 2012, 368 (2), 117–127.

42. Jonsson, T.; Stefansson, H.; Steinberg, S.; Jonsdottir, I.; Jonsson, P. V.; Snaedal, J.; Bjornsson, S.; Huttenlocher, J.; Levey, A. I.; Lah, J. J.; Rujescu, D.; Hampel, H.; Giegling, I.; Andreassen, O. A.; Engedal, K.; Ulstein, I.; Djurovic, S.; Ibrahim-Verbaas, C.; Hofman, A.; Ikram, M. A.; van Duijn, C. M.; Thorsteinsdottir, U.; Kong, A.; Stefansson, K., Variant of TREM2 associated with the risk of Alzheimer’s disease. The New England journal of medicine 2013, 368 (2), 107–16.

43. Ewers, M.; Franzmeier, N.; Suárez-Calvet, M.; Morenas-Rodriguez, E.; Caballero, M. A. A.; Kleinberger, G.; Piccio, L.; Cruchaga, C.; Deming, Y.; Dichgans, M.; Trojanowski, J. Q.; Shaw, L. M.; Weiner, M. W.; Haass, C., Increased soluble TREM2 in cerebrospinal fluid is associated with reduced cognitive and clinical decline in Alzheimer’s disease. Science translational medicine 2019, 11 (507).

44. Sampaolo, S.; Campos-Barros, A.; Mazziotti, G.; Carlomagno, S.; Sannino, V.; Amato, G.; Carella, C.; Di Iorio, G., Increased Cerebrospinal Fluid Levels of 3,3′,5′-Triiodothyronine in Patients with Alzheimer’s Disease. The Journal of Clinical Endocrinology & Metabolism 2005, 90 (1), 198–202.

45. Accorroni, A.; Giorgi, F. S.; Donzelli, R.; Lorenzini, L.; Prontera, C.; Saba, A.; Vergallo, A.; Tognoni, G.; Siciliano, G.; Baldacci, F.; Bonuccelli, U.; Clerico, A.; Zucchi, R., Thyroid hormone levels in the cerebrospinal fluid correlate with disease severity in euthyroid patients with Alzheimer’s disease. Endocrine 2017, 55 (3), 981–984.

46. Hogervorst, E.; Huppert, F.; Matthews, F. E.; Brayne, C., Thyroid function and cognitive decline in the MRC Cognitive Function and Ageing Study. Psychoneuroendocrinology 2008, 33 (7), 1013–22.

47. Davis, J. D.; Podolanczuk, A.; Donahue, J. E.; Stopa, E.; Hennessey, J. V.; Luo, L.-G.; Lim, Y.-P.; Stern, R. A., Thyroid hormone levels in the prefrontal cortex of post-mortem brains of Alzheimer’s disease patients. Curr Aging Sci 2008, 1 (3), 175–181.

48. Johansson, P.; Almqvist, E. G.; Johansson, J. O.; Mattsson, N.; Hansson, O.; Wallin, A.; Blennow, K.; Zetterberg, H.; Svensson, J., Reduced cerebrospinal fluid level of thyroxine in patients with Alzheimer’s disease. Psychoneuroendocrinology 2013, 38 (7), 1058–66.

49. Choi, H. J.; Byun, M. S.; Yi, D.; Sohn, B. K.; Lee, J. H.; Lee, J. Y.; Kim, Y. K.; Lee, D. Y., Associations of thyroid hormone serum levels with in-vivo Alzheimer’s disease pathologies. Alzheimer’s research & therapy 2017, 9 (1), 64.

50. Juárez-Cedillo, T.; Basurto-Acevedo, L.; Vega-García, S.; Sánchez-Rodríguez Martha, A.; Retana-Ugalde, R.; Juárez-Cedillo, E.; Gonzalez-Melendez Roberto, C.; Escobedo-de-la-Peña, J., Prevalence of thyroid dysfunction and its impact on cognition in older mexican adults: (SADEM study). Journal of endocrinological investigation 2017, 40 (9), 945–952.

51. Piccio, L.; Buonsanti, C.; Cella, M.; Tassi, I.; Schmidt, R. E.; Fenoglio, C.; Rinker, J., 2nd; Naismith, R. T.; Panina-Bordignon, P.; Passini, N.; Galimberti, D.; Scarpini, E.; Colonna, M.; Cross, A. H., Identification of soluble TREM-2 in the cerebrospinal fluid and its association with multiple sclerosis and CNS inflammation. Brain 2008, 131 (Pt 11), 3081–91.

52. Cignarella, F.; Filipello, F.; Bollman, B.; Cantoni, C.; Locca, A.; Mikesell, R.; Manis, M.; Ibrahim, A.; Deng, L.; Benitez, B. A.; Cruchaga, C.; Licastro, D.; Mihindukulasuriya, K.; Harari, O.; Buckland, M.; Holtzman, D. M.; Rosenthal, A.; Schwabe, T.; Tassi, I.; Piccio, L., TREM2 activation on microglia promotes myelin debris clearance and remyelination in a model of multiple sclerosis. Acta Neuropathologica 2020, 140 (4), 513–534.

53. Piccio, L.; Buonsanti, C.; Mariani, M.; Cella, M.; Gilfillan, S.; Cross, A. H.; Colonna, M.; Panina-Bordignon, P., Blockade of TREM-2 exacerbates experimental autoimmune encephalomyelitis. European journal of immunology 2007, 37 (5), 1290–301.

54. Jiang, T.; Tan, L.; Zhu, X.-C.; Zhang, Q.-Q.; Cao, L.; Tan, M.-S.; Gu, L.-Z.; Wang, H.-F.; Ding, Z.-Z.; Zhang, Y.-D.; Yu, J.-T., Upregulation of TREM2 Ameliorates Neuropathology and Rescues Spatial Cognitive Impairment in a Transgenic Mouse Model of Alzheimer’s Disease. Neuropsychopharmacology 2014, 39 (13), 2949–2962.

55. Lee, C. Y. D.; Daggett, A.; Gu, X.; Jiang, L. L.; Langfelder, P.; Li, X.; Wang, N.; Zhao, Y.; Park, C. S.; Cooper, Y.; Ferando, I.; Mody, I.; Coppola, G.; Xu, H.; Yang, X. W., Elevated TREM2 Gene Dosage Reprograms Microglia Responsivity and Ameliorates Pathological Phenotypes in Alzheimer’s Disease Models. Neuron 2018, 97 (5), 1032–1048.e5.

56. Takahashi, K.; Prinz, M.; Stagi, M.; Chechneva, O.; Neumann, H., TREM2-transduced myeloid precursors mediate nervous tissue debris clearance and facilitate recovery in an animal model of multiple sclerosis. PLoS medicine 2007, 4 (4), e124.

57. Perugorria, M. J.; Esparza-Baquer, A.; Oakley, F.; Labiano, I.; Korosec, A.; Jais, A.; Mann, J.; Tiniakos, D.; Santos-Laso, A.; Arbelaiz, A.; Gawish, R.; Sampedro, A.; Fontanellas, A.; Hijona, E.; Jimenez-Agüero, R.; Esterbauer, H.; Stoiber, D.; Bujanda, L.; Banales, J. M.; Knapp, S.; Sharif, O.; Mann, D. A., Non-parenchymal TREM-2 protects the liver from immune-mediated hepatocellular damage. Gut 2019, 68 (3), 533–546.

58. Mehta, P.; McAuley, D. F.; Brown, M.; Sanchez, E.; Tattersall, R. S.; Manson, J. J., COVID-19: consider cytokine storm syndromes and immunosuppression. The Lancet 2020, 395 (10229), 1033–1034.

59. Merad, M.; Martin, J. C., Pathological inflammation in patients with COVID-19: a key role for monocytes and macrophages. Nature Reviews Immunology 2020, 20 (6), 355–362.

60. Ferrara, S. J.; Scanlan, T. S., A CNS-Targeting Prodrug Strategy for Nuclear Receptor Modulators. Journal of Medicinal Chemistry 2020, 63 (17), 9742–9751.

61. Placzek, A. T.; Ferrara, S. J.; Hartley, M. D.; Sanford-Crane, H. S.; Meinig, J. M.; Scanlan, T. S., Sobetirome prodrug esters with enhanced blood–brain barrier permeability. Bioorganic & Medicinal Chemistry 2016, 24 (22), 5842–5854.

62. Placzek, A. T.; Scanlan, T. S., New synthetic routes to thyroid hormone analogs: d6-sobetirome, 3H-sobetirome, and the antagonist NH-3. Tetrahedron 2015, 71 (35), 5946–5951.

63. Hackenmueller, S. A.; Marchini, M.; Saba, A.; Zucchi, R.; Scanlan, T. S., Biosynthesis of 3-Iodothyronamine (T1AM) Is Dependent on the Sodium-Iodide Symporter and Thyroperoxidase but Does Not Involve Extrathyroidal Metabolism of T4. Endocrinology 2012, 153 (11), 5659–5667.

64. Pfaffl, M. W., A new mathematical model for relative quantification in real-time RT–PCR. Nucleic Acids Research 2001, 29 (9), e45–e45.

65. Ferrara, S. J.; Bourdette, D.; Scanlan, T. S., Hypothalamic-Pituitary-Thyroid Axis Perturbations in Male Mice by CNS-Penetrating Thyromimetics. Endocrinology 2018, 159 (7), 2733–2740.

66. Humbert-Claude, M.; Duc, D.; Dwir, D.; Thieren, L.; Sandström von Tobel, J.; Begka, C.; Legueux, F.; Velin, D.; Maillard, M. H.; Do, K. Q.; Monnet-Tschudi, F.; Tenenbaum, L., Tollip, an early regulator of the acute inflammatory response in the substantia nigra. Journal of Neuroinflammation 2016, 13 (1), 303.

67. Sun, Y.; Ma, J.; Li, D.; Li, P.; Zhou, X.; Li, Y.; He, Z.; Qin, L.; Liang, L.; Luo, X., Interleukin-10 inhibits interleukin-1β production and inflammasome activation of microglia in epileptic seizures. Journal of Neuroinflammation 2019, 16 (1), 66.

68. Zhao, H.; Liu, A.; Shen, L.; Xu, C.; Zhu, Z.; Yang, J.; Han, X.; Bao, F.; Yang, W., Isoforskolin downregulates proinflammatory responses induced by Borrelia burgdorferi basic membrane protein A. Exp Ther Med 2017, 14 (6), 5974–5980.

69. Min, Z.; Tang, Y.; Hu, X.-T.; Zhu, B.-L.; Ma, Y.-L.; Zha, J.-S.; Deng, X.-J.; Yan, Z.; Chen, G.-J., Cosmosiin Increases ADAM10 Expression via Mechanisms Involving 5’UTR and PI3K Signaling. Frontiers in Molecular Neuroscience 2018, 11 (198).

70. Rai, V.; Rao, V. H.; Shao, Z.; Agrawal, D. K., Dendritic Cells Expressing Triggering Receptor Expressed on Myeloid Cells-1 Correlate with Plaque Stability in Symptomatic and Asymptomatic Patients with Carotid Stenosis. PLOS ONE 2016, 11 (5), e0154802.

