## Supporting Information for "*TREM2* is thyroid hormone regulated making the TREM2 pathway druggable with ligands for thyroid hormone receptor"

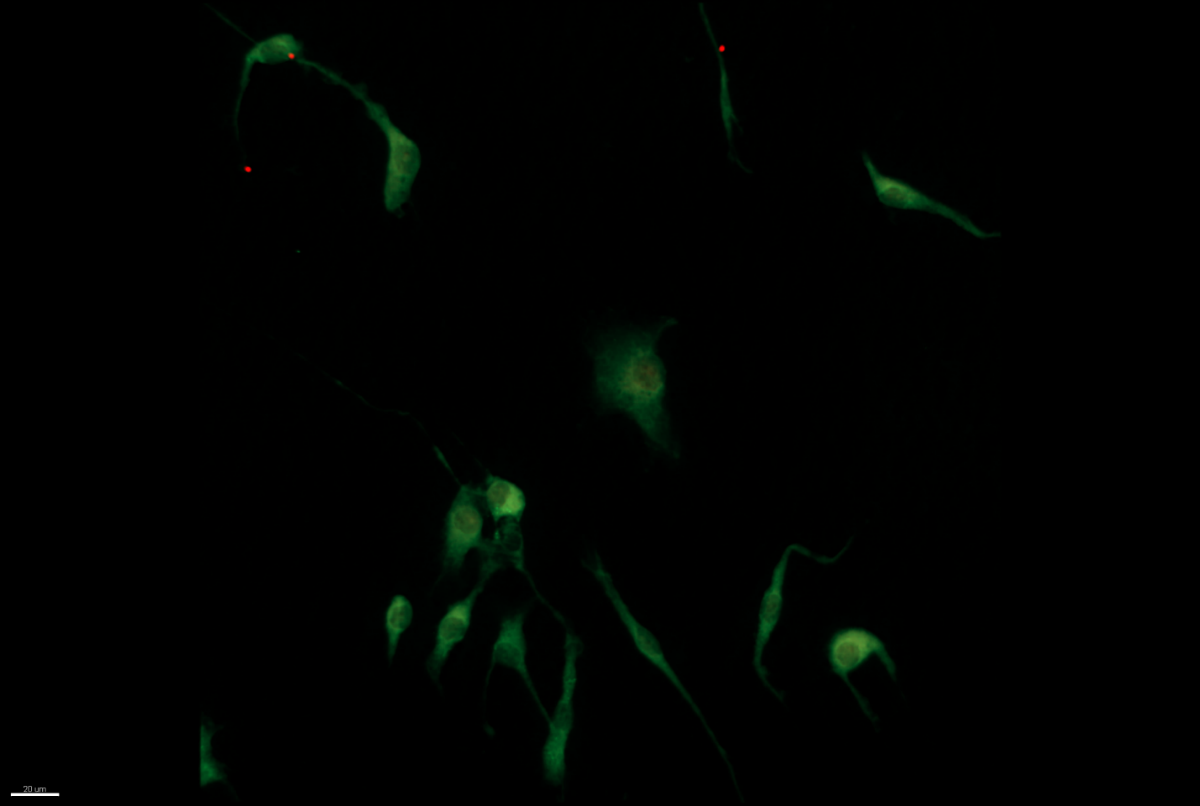

**Supplemental Figure S1**: Full-field blown-up image of the DMSO vehicle-treated mouse primary microglia culture representative field in Fig. 4a.

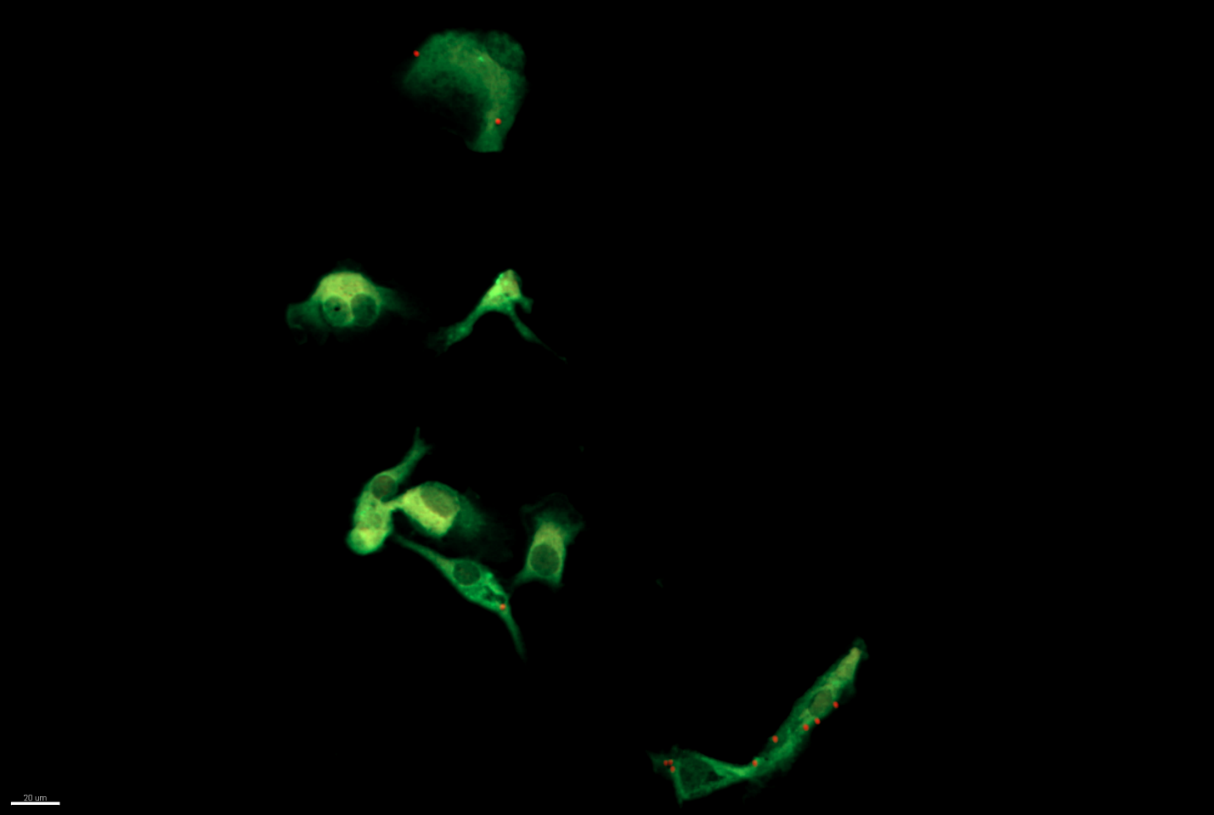

**Supplemental Figure S2**: Full-field blown-up image of the 10 nM T3-treated mouse primary microglia culture representative field in Fig. 4b.

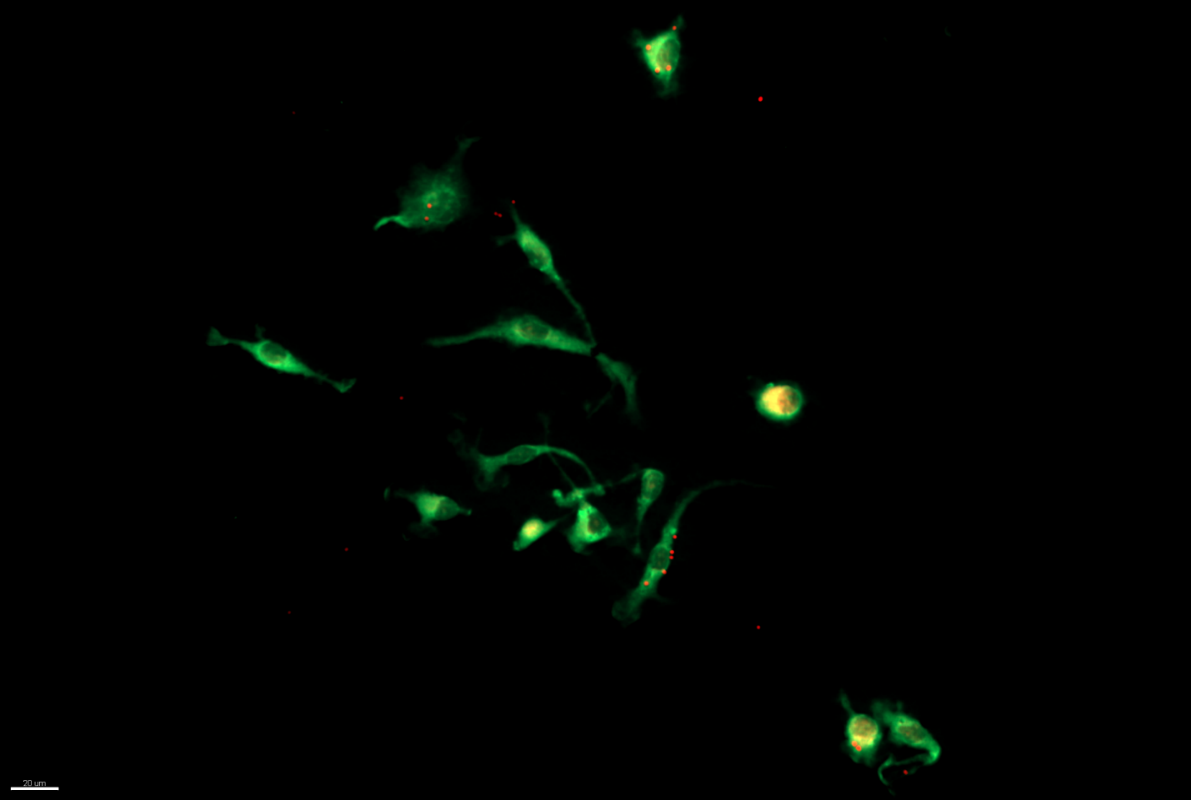

**Supplemental Figure S3**: Full-field blown-up image of the 1μM sobetirome-treated mouse primary microglia culture representative field in Fig. 4c.

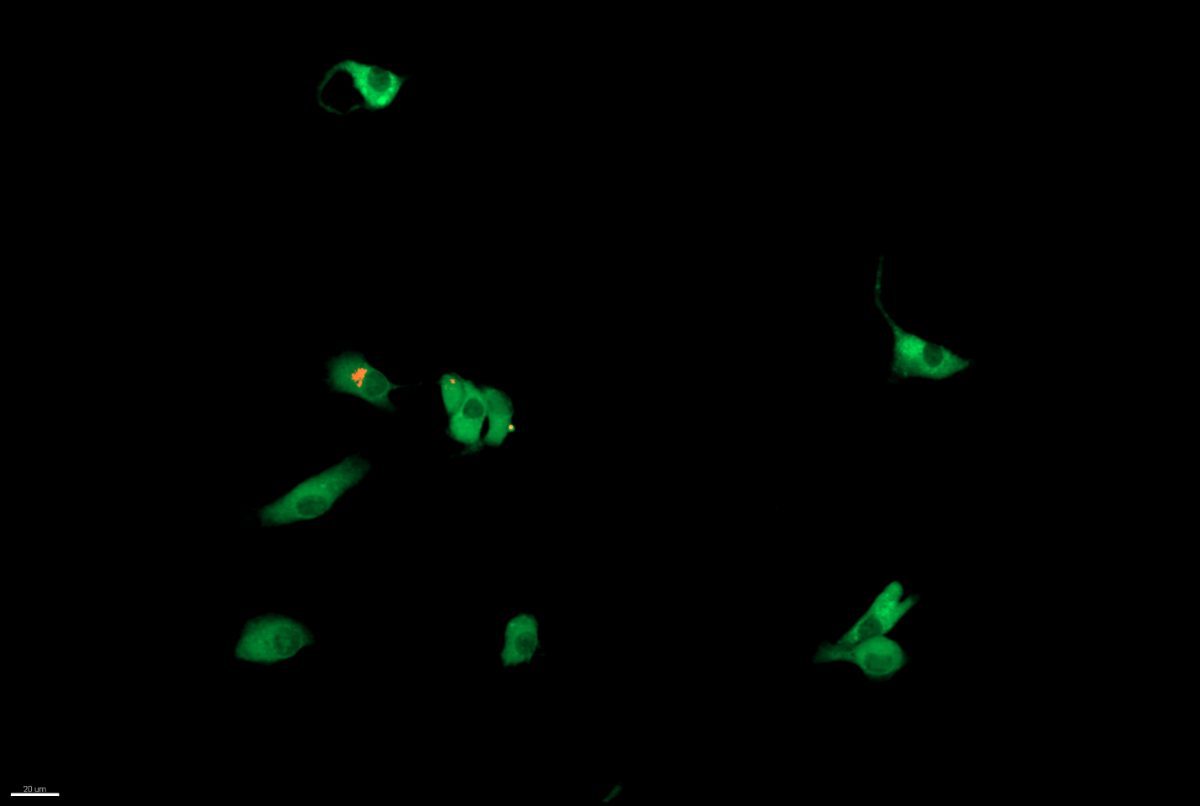

**Supplemental Figure S4**: Full-field blown-up image of the 2μM NH-3-treated mouse primary microglia culture representative field in Fig. 4d. Note: Aberrant autofluorescence was observed in the indicated cell (white arrow) which is not due to the presence of bead(s).

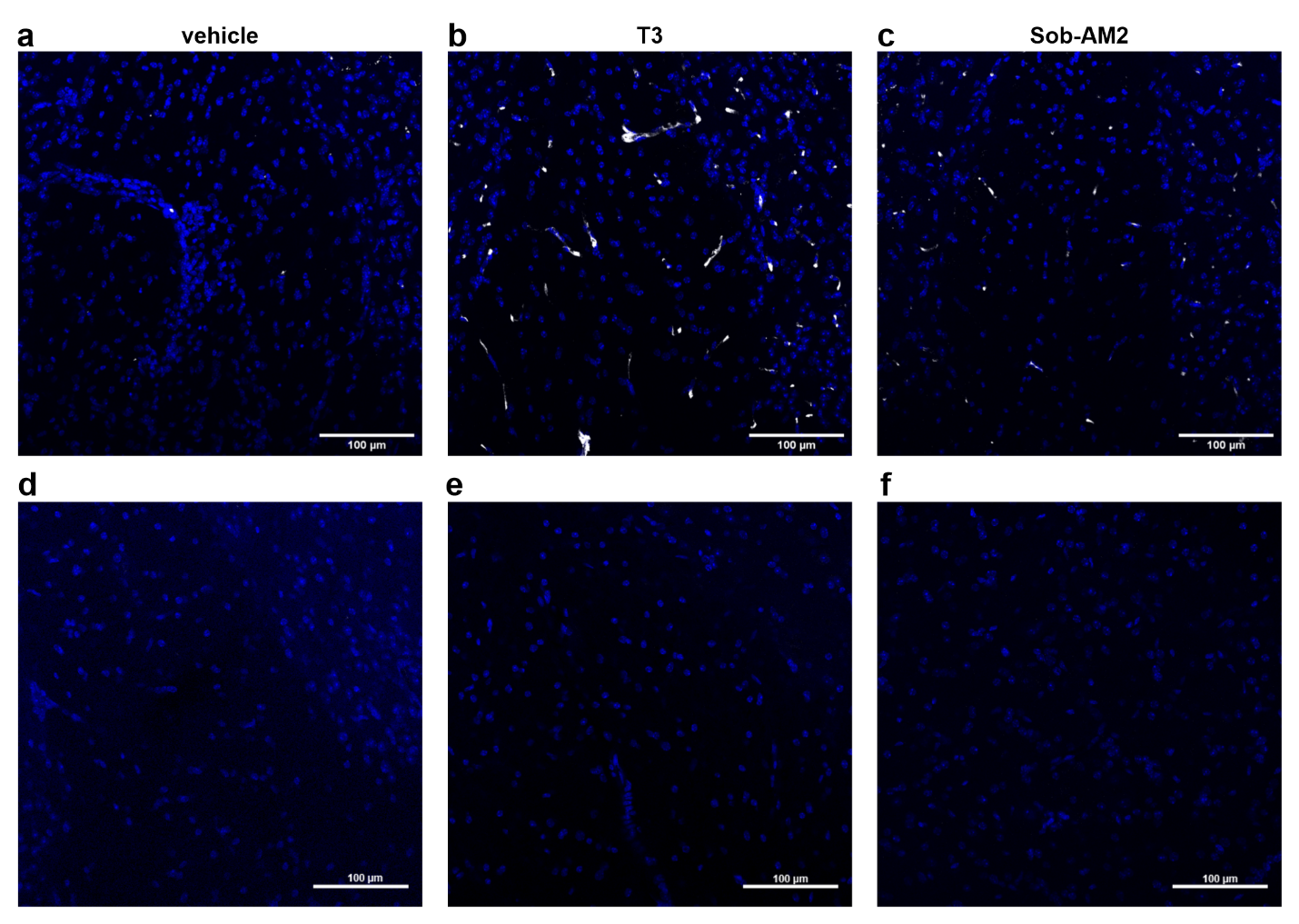

**Supplemental Figure S5**: **(a-c)** DAPI and TREM2 co-staining for the representative images in Fig. 5. **(d-f)** No primary controls for each group showing DAPI and TREM2 channels. Scale bars: 100 μm.

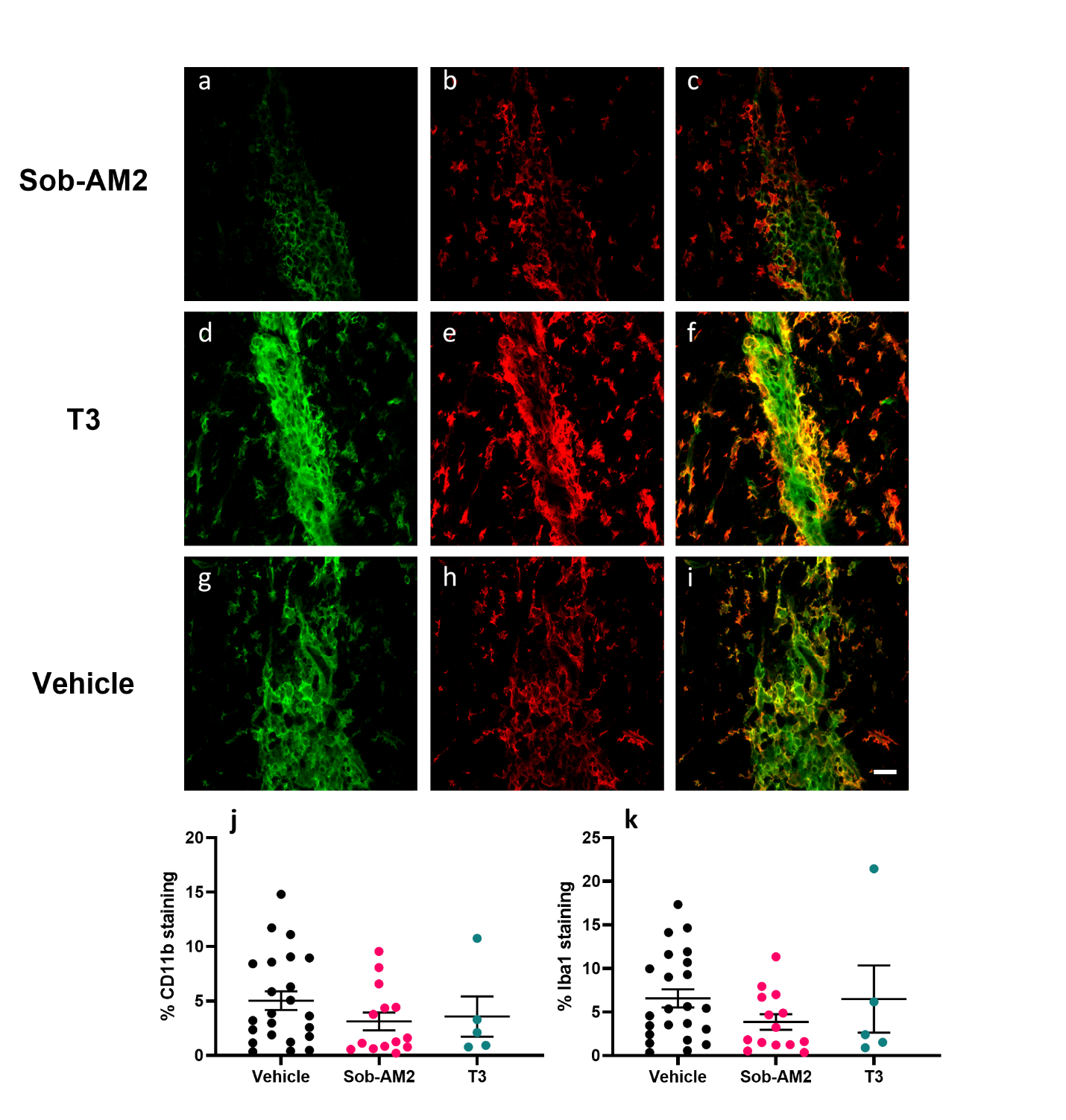

**Supplemental Figure S6:** Representative CD11b (green) and Iba1 (red) staining of dorsal spinal cord sections from Sob-AM2 (**a-c**,1 mg/kg), T3 (**d-f**, 0.4 mg/kg), and vehicle (**g-i**, DMSO and NaOH) treated mice. Quantification of percent CD11b staining (**j**) and Iba1 staining (**k**) from each representative group. There is no statistically significant difference between any group in **j** and **k** as determined by one-way ANOVA (P < 0.05) and Mann-Whitney U tests between vehicle and specific group. Scale bar is 20 μm.
